# Nutrient limitation magnifies fitness costs of antimalarial drug resistance mutations

**DOI:** 10.1101/2021.08.02.454586

**Authors:** Shalini Nair, Xue Li, Grace A. Arya, Marina McDew-White, Marco Ferrari, Tim Anderson

**Affiliations:** Disease Intervention and Prevention Program, Texas Biomedical Research Institute, San Antonio, Texas 78245

**Keywords:** Resource limitation, Plasmodium falciparum, Artemisinin resistance, fitness costs, compensation

## Abstract

Drug resistance mutations tend to disrupt key physiological processes, and therefore carry a fitness cost. The size of these fitness costs is a central determinant of the rate of spread of these mutations in natural populations so are important to quantify. Head-to-head competition assays provide a standard approach to measuring differential fitness, and have been used extensively for malaria parasites. These assays typically use standardized culture media, containing RPMI 1640, which has a 1.4 to 5.5-fold (mean: 2.6-fold) higher concentration of amino acids than human blood. In this rich media we predict that fitness costs will be underestimated because resource competition is weak. We tested this prediction using an artemisinin sensitive parasite edited to contain *kelch-C580Y* or *R561H* mutations conferring resistance to artemisinin or synonymous control mutations. We examined the impact of these single amino acid mutations on fitness, using replicated head-to head competition experiments conducted in media containing (i) normal RPMI, (ii) modified RPMI with reduced amino acid concentration, (iii) RPMI containing only isoleucine, or (iv) 3-fold diluted RPMI. We found a significant 1.3 to 1.4-fold increase in fitness costs measured in modified and isoleucine-only media relative to normal media, while fitness costs were 2.5-fold higher in diluted media. We conclude that fitness costs are strongly affected by media composition and will be significantly underestimated in normal RPMI. Elevated fitness costs in nature will limit spread of ART-resistance but will also promote evolution of compensatory mutations that restore fitness, and can be exploited to maximize selection in laboratory experiments.

## INTRODUCTION

Drug resistance mutations typically carry a fitness cost: pathogens carrying these mutations tend to be outcompeted by wild type pathogens when they are in direct competition, but have a fitness advantage over wildtype parasites when exposed to drug treatment (1, 2). Such fitness costs occur because drug resistance mutations interfere with key metabolic pathways – for example, in the case of the malaria parasite *Plasmodium falciparum*, mutations conferring resistance to chloroquine in *pfcrt* reduces peptide transport capacity (3), kelch13 mutations underlying artemisinin resistance reduced levels of endocytosis (4), and dihydrofolate reductase mutations reduce efficiency of folate synthesis (5, 6). Fitness costs associated with chloroquine resistance provide an explanation for the rapid disappearance of drug resistance (*pfcrt*) alleles from Malawi and other countries after removal of chloroquine as first line treatment (7, 8). Fitness costs may also help explain why resistance alleles have historically evolved in low transmission regions such as SE Asia and S. America, rather than in sub-Saharan Africa where most parasites are found (9). In such low transmission regions most infections contain single parasite genotypes (10), so direct competition between drug resistant and drug sensitive parasites is minimized allowing emergence and transmission of drug resistant mutations to new hosts (11). Fitness costs are also an important driver of compensatory changes that ameliorate fitness (12, 13): if costs are high then there is strong selection for compensatory changes, while if costs are low, such compensatory changes are less likely to emerge (2, 14).

Fitness costs are now widely measured in the laboratory in studies of antimalarial drug resistance (15–19). This is typically done using competition experiments in which resistant and sensitive parasites are competed in culture, and their proportions are monitored over time to calculate selection coefficients which measures the relative fitness advantage each generation (19). For example, parasite clones with a selection coefficient of -0.1 produced 10% less offspring each generation than the competing parasite. A concern is that the magnitude of fitness costs determined using competition-based assays are likely to be dependent on the media used. Limitation of critical resources (e.g. nutrients, space, light) determines the strength of competition between organisms (20). Therefore, the strength of competition and size of fitness costs measured is expected to depend on the degree of resource limitation (21, 22). There is strong precedent for the role of nutrient limitation in mediating bacterial competition (23–25). Furthermore, comparisons of non-isogenic artemisinin-resistant and sensitive parasites from Cambodia, have suggested that fitness costs are exacerbated in conductions of amino acid starvation (26).

RPMI 1640, a nutrient rich media designed to maximize growth of multiple cell types, forms the basis of malaria culture media. Red blood cells, and a lipid source – either human serum or a bovine serum substitute (Albumax) – are then added to RPMI to make complete media. The resultant complete media contains much higher levels of many nutrients than human blood (27–29). For example, amino acids are present at concentrations 1.4-5.5-fold (mean: 2.6-fold) greater than observed in human blood. Of these amino acids, isoleucine is critical for malaria parasite growth (30), because other amino acids can be obtained from hemoglobin digestion (31). However, isoleucine is found at 5.5-fold greater concentration in RPMI than in human blood (Table 1) (27, 28).

**Table 1.** Nutrient level in RPMI used to construct four different growth media used in this study. Modified and Starvation RPMI were made to order, while diluted RPMI was mixed with Hank’s balance salt solution. Nutrient levels found in human plasma are shown for reference. Only the 7 amino acids found in normal RPMI 1640 are shown here. Table S1 shows a full list of aminio acids and other compounds that are found in human plasma, but not in normal or altered RPMI 1640.

| Nutrient | Standard RPMI | Modified RPMI | Starvation RPMI | Diluted RPMI (3-fold HBSS<sup>a</sup>) | Human Plasma<sup>b</sup> |
| --- | --- | --- | --- | --- | --- |
| L-Glutamate<sup>1</sup> | 0.136 | 0.053 | -- | 0.045 | 0.080 |
| L-Glutamine<sup>5</sup> | 2.055 | 0.611 | -- | 0.685 | 0.550 |
| L-Isoleucine<sup>1c</sup> | 0.382 | 0.074 | 0.074 | 0.127 | 0.070 |
| L-Methionine<sup>1</sup> | 0.101 | 0.030 | -- | 0.034 | 0.030 |
| L-Proline<sup>1</sup> | 0.174 | 0.204 | -- | 0.058 | 0.200 |
| L-Tyrosine<sup>1</sup> | 0.111 | 0.067 | -- | 0.037 | 0.080 |
| L-Cystine<sup>1</sup> | 0.208 | 0.114 | -- | 0.069 | 0.100 |
| **Vitamins** |  |  |  |  |  |
| Biotin | 0.001 | 0.001 | 0.001 | 2.70E-04 | 0.001 |
| Choline chloride | 0.021 | 0.021 | 0.021 | 0.007 | 0.022 |
| D-Calcium pantothenate | 0.001 | 0.001 | 0.001 | 1.70E-04 | 0.001 |
| Folic Acid | 0.002 | 0.002 | 0.002 | 7.50E-04 | 0.002 |
| Niacinamide | 0.008 | 0.008 | 0.008 | 0.003 | 0.008 |
| Para-Aminobenzoic Acid<sup>d</sup> | 0.007 | 0.007 | 0.007 | 0.002 | 0.007 |
| Pyridoxine hydrochloride | 0.005 | 0.005 | 0.005 | 0.002 | 0.005 |
| Riboflavin | 0.001 | 0.001 | 0.001 | 1.70E-04 | 0.001 |
| Thiamine hydrochloride | 0.003 | 0.003 | 0.003 | 9.80E-04 | 0.003 |
| Vitamin B12 | 3.76E-06 | 3.76E-06 | 3.76E-06 | 1.25E-06 | 3.70E-06 |
| D-Pantothenic acid | 0.001 | 2.30E-04 | 2.30E-04 | 1.70E-04 | 0.001 |
| i-Inositol | 0.194 | 0.194 | 0.194 | 0.065 | 0.194 |
| **Inorganic Salts** |  |  |  |  |  |
| Calcium nitrate (Ca(NO<sub>3</sub>)<sub>2</sub>) | 0.424 | 0.424 | 0.424 | 0.141 | 0.040 |
| Magnesium Sulfate (MgSO<sub>4</sub>) | 0.407 | 0.407 | 0.407 | 0.407 | 0.350 |
| Potassium Chloride (KCl) | 5.333 | 5.333 | 5.333 | 5.333 | 4.100 |
| Sodium Bicarbonate (NaHCO<sub>3</sub>) | 23.810 | 23.810 | 23.810 | 10.714 | 24.000 |
| Sodium Chloride (NaCl) | 103.448 | 103.448 | 103.448 | 126.437 | 105.000 |
| Sodium Phosphate dibasic | 5.634 | 5.634 | 5.634 | 2.103 | -- |
| **Other Components** |  |  |  |  |  |
| D-Glucose (Dextrose) | 11.111 | 5.500 | 5.500 | 7.370 | 5.000 |
| Glutathione (reduced) | 0.003 | 2.30E-04 | 2.30E-04 | 0.001 | 0.025 |

- HBSS (Hanks balanced salt solution) - Human plasma values are from Desai et al. (28) - Isoleucine is required by *P. falciparum* and cannot be synthesized from breakdown of hemoglobin - Para aminobenzoic acid levels limit growth of rodent malaria parasites

These experiments were designed to investigate the impact of media composition on measurement of fitness costs of antimalarial drug resistance, with a focus on amino acid composition. We expect that the strength of competition between parasites in competitive growth assays will depend on resource limitation: competition will be greatest when resources such as isoleucine are limiting, while when resources are abundant competition will be minimized. We also expect that metabolically impaired drug resistant parasites will be disproportionately affected compared with wildtype parasites when nutrients are limiting, but will suffer minimal costs when nutrients are plentiful. We predicted that fitness costs associated with drug resistance will be underestimated when measured in media containing normal RPMI, compared with media containing amino levels more comparable to that found in human blood.

## MATERIALS AND METHODS

### Experimental design

Fig 1 shows an overview of the experimental design.

**Figure 1.**
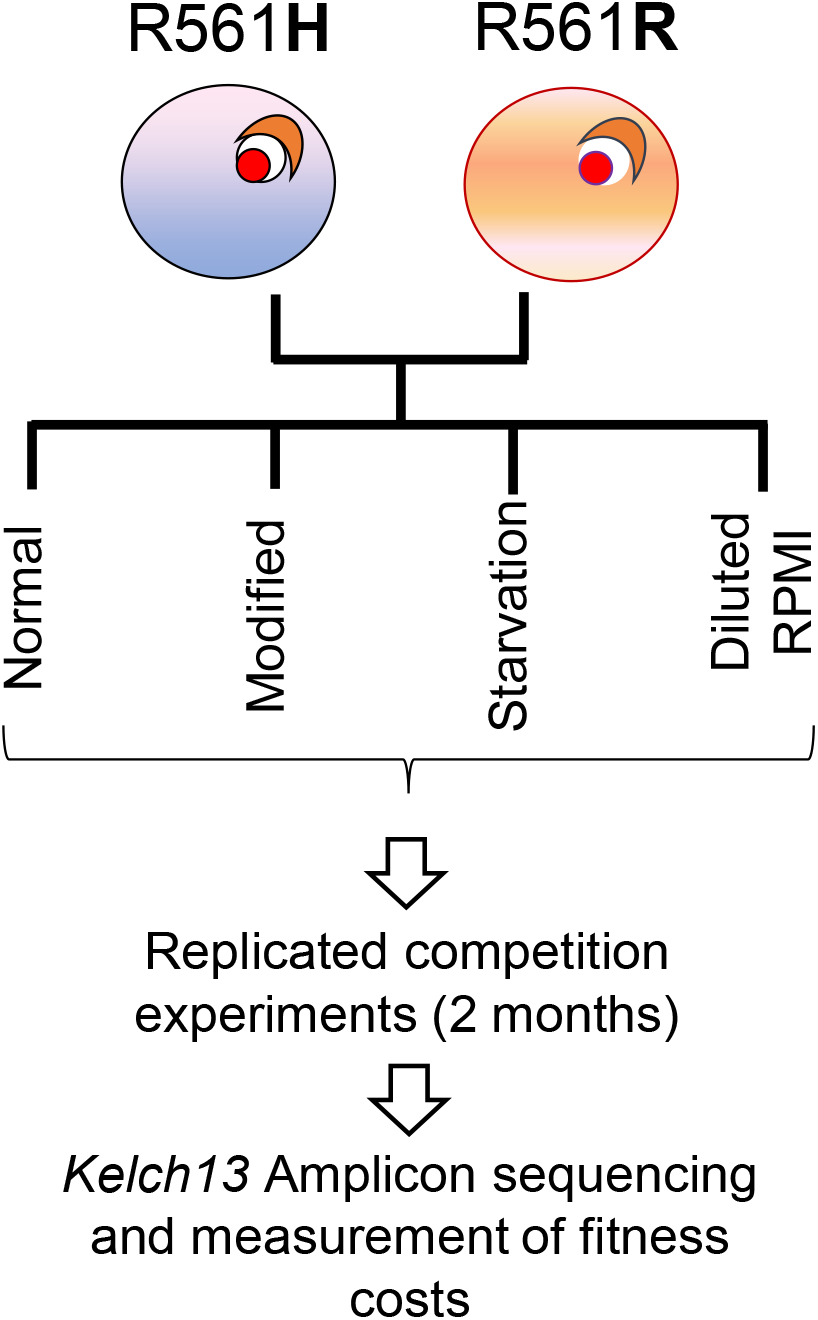
Overview of Project design. CRISPR modified parasites with either wild type or kelch13-R561H were mixed in 50:50 proportions, and replicated competition experiments were conducted in 4 different media types over 2 months. We used amplicon sequencing to determine frequencies of competing parasites and determine selection coefficients. There experiments were also conducted with wild type and C580Y.

### Parasites

We used four parasite clones in this work: NHP4302^C580Y^, NHP4302^C580C^, NHP4302^R561H^, and NHP4302^R561R^. These were all generated using CRISPR/Cas9 editing by introducing non-synonymous substitutions of key amino acids determining ART-resistance (NHP4302^C580Y^ and NHP4302^R561H^) or control synonymous mutations in the same codons NHP4302^C580C^ and NHP4302^R561R^) (19). The original parasite used for editing (NHP4302) was cloned from a parasite isolate recovered from a patient visiting the Wang Pha clinic run by the Shoklo Malaria Research Unit (SMRU) in 2008. This isolate had a wild-type kelch13 allele and cleared rapidly from the infected patients blood following treatment (clearance rate half-life (*T_1/2_P*) = 1.98 h) (19).

### Media composition

These experiments measured the fitness costs of isogenic parasites with or without drug resistance mutations in the kelch13 locus, using media that differed in composition. We cultured asexual blood-stage parasites in culture medium (RPMI 1640 supplemented with 2 mM L-glutamine, 25 mM HEPES, and 50 g/liter gentamicin, 0.1M hypoxanthine, and 0.4% albumax as a serum source) at 2% hematocrit and maintained them at 37°C with 5% O2, 5% CO2, and 90% N2). However, the composition of the RPMI differed in these experiments (Table 1). In brief, the four media types used were: (i) <u>Normal RPMI</u> 1640 (ii) <u>Modified RPMI</u> contained reduced levels of amino acids that approximates that found in human blood (28) (iii) <u>Starvation RPMI</u> contained isoleucine only at the same levels as in modified media. Isoleucine is the only essential amino acid for *P. falciparum* growth all others can be obtained from hemoglobin digestion (30, 31) (iv) <u>Diluted RPMI</u> contained a 3-fold dilution of RPMI in Hank’s basal salt solution (HBSS). HBSS contains salts at the same concentration as RPMI 1640, but lacks amino acids and trace elements found in RPMI. Red blood cells from different donors may have a large influence on *P. falciparum* growth rates (32): we therefore used a single donor (O+) for all experiments.

### Competition experiments

We conducted two sets of competition experiments (i) NHP4302^C580Y^ (ART-R) against NHP4302^C580C^ (ART-S) (ii) NHP4302^R561H^ (ART-R) against NHP4302^R561R^ (ART-S). These experiments were conducted in the four media types, with 6 replicates per experiment. In brief, cultures of the two parasite lines for comparison were (i) synchronized to 80% late schizonts using MACS purification columns (Miltenyi Biotec) (ii) diluted to 0.1% density and grown overnight to generate ring stage parasites (iii) mixed in equal proportions at a density of 0.1%. We ran 6 replicates of each competition experiment in the wells of a six well plate. These were then maintained for 2 months, with removal of aliquots (80 μl packed red cells) and dilution down to 0.1% from each culture at four-day intervals.

### Selection coefficient estimation

We used amplicon sequencing to determine frequencies of the two competing parasites in each experiment (19). In brief, we amplified a 249 bp region of kelch13 locus from each sampling time point of the competition experiments. These were then illumina sequenced (2 × 250bp reads) to high read depth on a MiSeq. We used oligos with individual barcodes to allow pooling of PCR products from multiple different reactions for sequencing (Read1 seq primer: TATGGTAATTGTACGTCAAATGGTAGAATTTATTGTATTGGGGGATATGATGGC; Read2 seq primer: AGTCAGCCAGCCAGGCATATGGAAATTGTTC; Index sequence primer: AATGATACGGCGACCACCGAGATCTACACGCT). We counted the number of reads from the two competing parasite lines to determine their frequency within each culture. We then plotted the natural log (proportion of resistant alleles) over time in asexual cycle (1 asexual generation = 48 hours) and measured the slope of the best fit line to give the selection coefficient. Scripts used are provided in github (https://github.com/emilyli0325/Amplicon.analysis.git): these have been updated from those used in (19). We remove data that only contains reads from one allele (allele frequency equals 0 or 1) and all data after those points; we use Cook’s Distance (33) to detect outliers that don’t fit the linear regression with a cut-off of 4/n; the first remaining data point was then defined as starting time point – asexual cycle 0.

### Statistical analysis

We analyzed the slopes from the 6 replicate cultures using the R program metaphor to conduct a random effects analysis, to determine the impact of media type on fitness costs.

## RESULTS

The final dataset comprised fitness cost measures for the R561H mutation in 4 different media types and for the C580Y mutation in 3 different media types (we did not conduct competition using 3-fold diluted RPMI for this mutation). Each of these 7 competition assays had 6 replicates, with 15 aliquots collected at four day intervals over two months. The amplicon sequence data used to quantify frequencies of the competing parasites is summarized in Table S1. We calculated allele frequencies from 5885±3421 reads per sampling time point, after excluding time points with < 100 reads.

We found a significant impact of media composition on fitness costs for both R561H and C580Y (Table 2, Figure 2). Fitness costs were lowest in media containing normal RPMI, but were 1.3-fold higher in both modified (t-test, p = 0.0192) or starvation media (t-test, p = 0.0148) for C580Y. We observed parallel changes for R561H, with a 1.38-fold increase in fitness costs in modified media (t-test, p = 9.01 × 10^−04^) and a 1.5 fold increase in starvation media (t-test, 3.10 × 10^-04^) (Figure 3). We did not see any significant difference between modified and starvation media for either R561H (t-test, p=0.89) or C580Y (t-test, p=0.97).

**Table 2.**
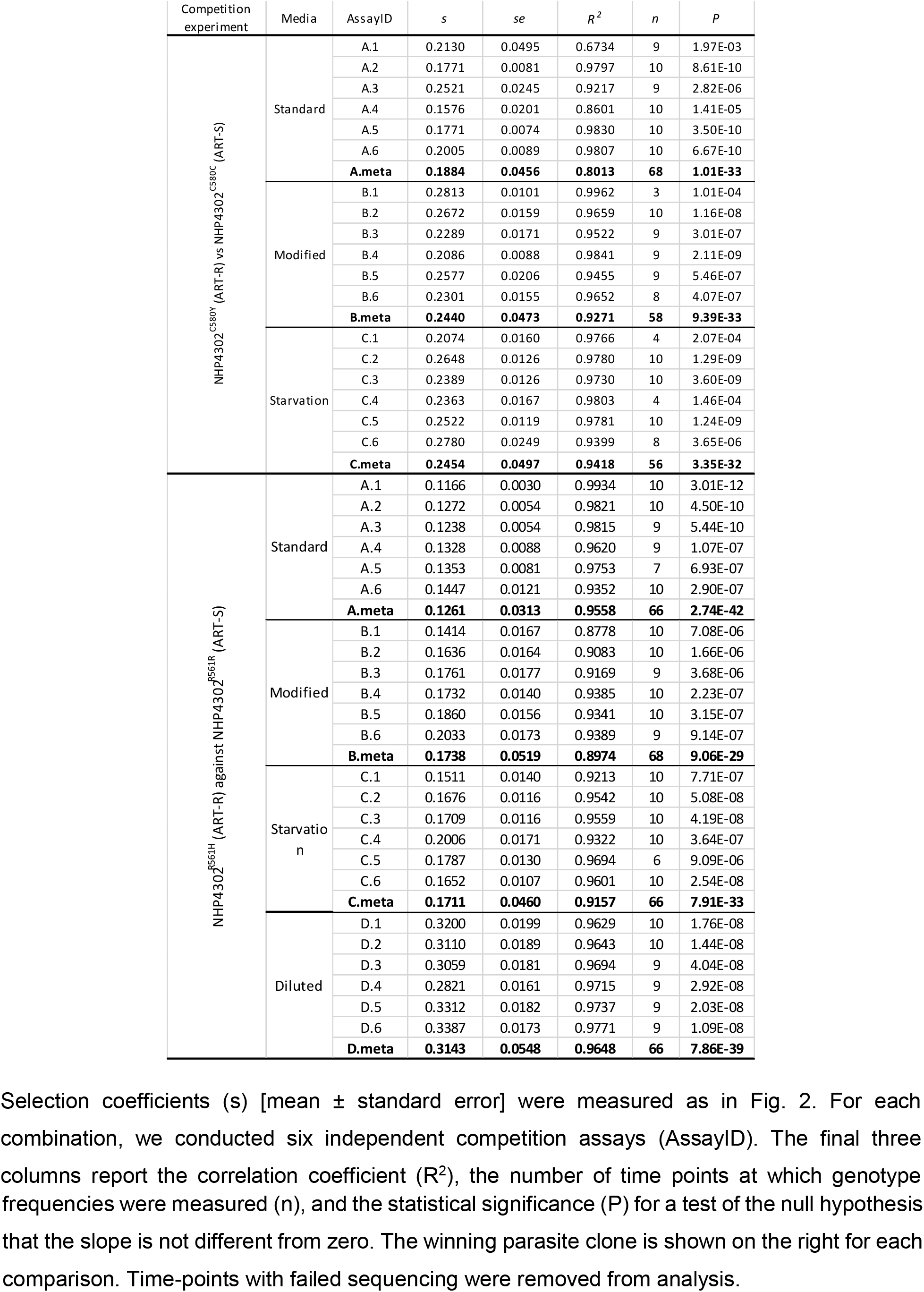
Calculation and summary for selection coefficients of all competition experiments.

**Table 3.** Pairwise comparison of fitness costs (s) of drug resistance mutations measured in different media. A. C580Y, B. R561H. Upper triangle shows significance of t-tests comparing fitness cost measured in different media types. Grey shading, p< 0.05; black shading/white text, p<0.001. Lower triangle shows fold-change in fitness costs measured in different media. Results are summarized graphically in figure 2, while all replicates are compared statistically in Table 2.

**A. C580Y**
|  | Standard | Modified | Starvation | Diluted |
| --- | --- | --- | --- | --- |
| Standard | -- | 0.019 | 0.015 | nd |
| Modified | 1.295 | -- | 0.967 | nd |
| Starvation | 1.303 | 1.006 | -- | nd |
| Diluted | nd | nd | nd | -- |

**B. R561H**
|  | Standard | Modified | Starvation | Diluted |
| --- | --- | --- | --- | --- |
| Standard | -- | 9.01E-04 | 3.10E-04 | 1.94E-09 |
| Modified | 1.38 | -- | 0.89 | 3.18E-07 |
| Starvation | 1.36 | 0.98 | -- | 1.04E-07 |
| Diluted | 2.49 | 1.81 | 1.84 | -- |

**Figure 2.**
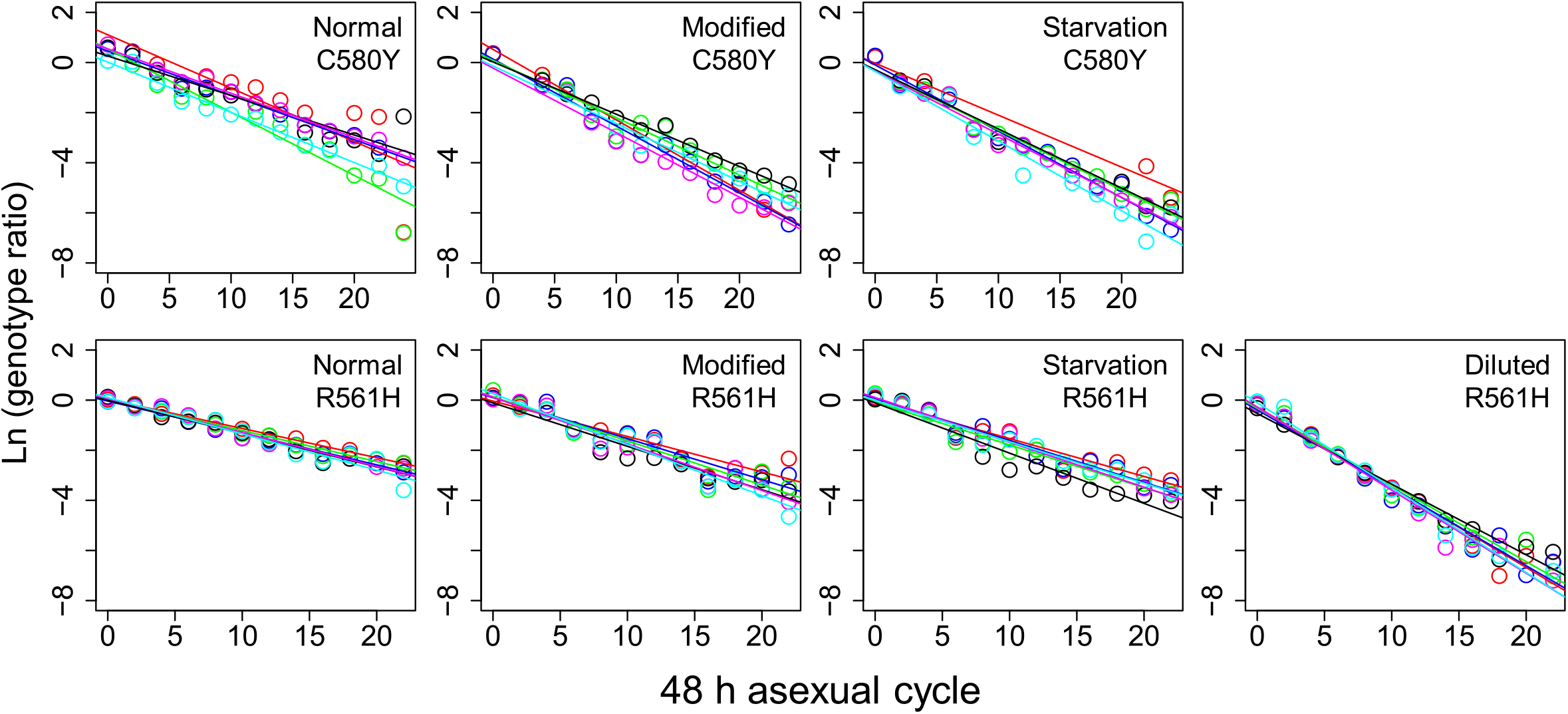
Trajectory of competing parasites observed in replicate experiments. The plots show the natural log of the parasite ratio against time between competing parasites. The slope for the least-squares fit provides an estimate of the selection coefficient (*s*). Each color represents one replicate.

**Figure 3.**
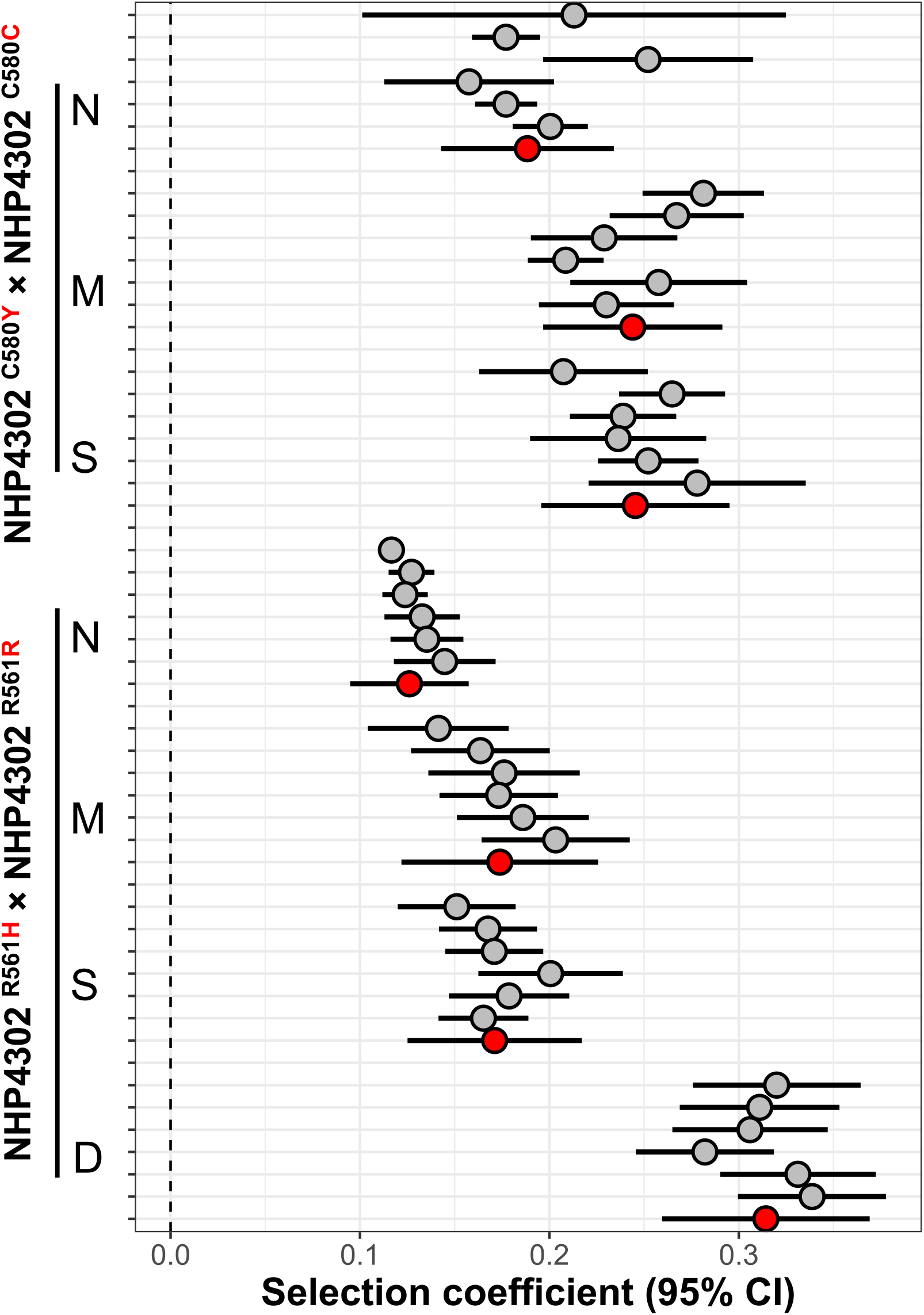
Outcome of competition experiments in different media. The plot shows selection coefficients (*s*) with 95% confidence intervals for competition experiments conducted in different media. Six replicate competition experiments were conducted for each set: Grey points shows *s* for each replicate, while the red points show meta-analysis results for each experimental comparison. The “winning” parasite is shown on the right for each comparison.

We found the largest fitness cost differences between normal RPMI and 3-fold diluted RPMI. In this media R561H showed a fitness cost of 0.31±0.05 compared with 0.13±0.03 in normal RPMI (t-test, p=1.94 × 10^-09^). This represents a 2.49-fold difference in fitness costs depending on media composition used (Figure 3).

## DISCUSSION

### Nutrient limitation increases fitness costs in *P. falciparum*

These experiments were designed to test the prediction that fitness costs are dependent on the nutrient composition of parasite growth media. We observed a modest (1.29 - 1.38-fold), but significant, increase in selection costs when parasites were grown in media containing reduced amino acid composition designed to mimic that found in human blood. This was observed for both C580Y and R561H mutations providing strong support for this hypothesis.

Malaria parasites are able to synthesize all amino acids except for isoleucine (30, 31), hence we would expect that isoleucine concentrations might be of particular importance. However, as artemisinin resistant parasites show reduced endocytosis of hemoglobin compared to wildtype parasites (4), access to amino acids synthesized from products of hemoglobin digestion may be also limiting in ART-R parasites. Interestingly, we found that starvation media, containing isoleucine only in concentrations mimicking normal blood, also resulted in comparable increases in fitness costs to that observed in modified media containing 7 amino acids. These results are consistent with isoleucine levels being sufficient to cause the observed changes in fitness costs.

Three-fold diluted RPMI contains slightly higher isoleucine concentrations than the modified and starvation media (Table 1), so would be expected to have a less pronounced impact on fitness measures if isoleucine is solely responsible. However, diluted media had a massive (2.5-fold) impact on fitness coefficients, suggesting that limitation of other RPMI components impacts fitness coefficients. We used Hanks balanced salt solution as a diluent for RPMI. This matches the salt concentration of RPMI1640 but lacks several trace elements. We infer that one of more of these trace elements are critical for *Plasmodium* growth. A likely candidate is para-aminobenzoic acid (pABA), which is required for folate biosynthesis. As folate is needed to for pyrimidine synthesis and methionine metabolism, pABA is critical for malaria parasite growth (34, 35). Previous studies have shown that pABA limits malaria parasite growth rates (35). Furthermore, manipulation of pABA levels alters the outcome of competition between pyrimethamine resistant and sensitive malaria parasites (*P. chabaudi*) in rodent malaria parasites, providing an in vivo demonstration of the importance of nutrient limitation in determined competitive outcomes (21, 22).

Bunditvorapoom et al (26) observed that differences in fitness were exacerbated in low amino acid conditions in comparisons between two resistant and two sensitive parasites isolated from Cambodian patients. Ring stage ART-S – but not ART-R – parasites, were able to mature to trophozoites under low amino acid conditions, while both ART-S and ART-R parasites made the ring -> trophozoite transition under standard culture conductions. Our results confirm and extend this interesting observation, by quantifying the impact of amino acid starvation on fitness in isogenic CRISPR-edited parasites.

Fitness costs are widely measured in the laboratory to understand why particular resistance mutations are spreading in nature. These studies are particularly powerful when combined with CRISPR, because single mutations can be examined on the same genetic background (16–19). An important point is that fitness costs are likely to be underestimated by studies using normal RPMI, because this is a rich media designed to allow growth of multiple cell types or microbe species. We can obtain more accurate estimates of relative fitness using RPMI with reduced amino acid levels. However, we note that multiple aspects of media used in malaria culture (lipid source, immunity, movement) differ from that observed in human blood (27–29), so translating fitness cost measured in the laboratory to the situation within patients is problematic.

### Practical consequences of resource limitation

The dramatic effects of resource limitation on competitive interactions between parasites can be effectively harnessed for experimental work. We provide three examples:

Measuring fitness costs in malaria parasites typically requires 1-2 months of co-culture to resolve competitive outcomes. Resource limitation allows significant shortening of these experiments, because fitness costs are magnified, and one of the two competitors will reach fixation. For example, in our experiments using the *kelch-R561H* resistance mutation conferring artemisinin resistance, parasites bearing the NHP4302^R561H^ allele were still present on day 50 in normal media, but were eliminated by day 26 in diluted media. Hence, we can shorten these experiments and better resolve small differences in fitness.

We have recently utilized malaria parasite genetic crosses to generate large progeny pools, which can then be monitored over time to examine relative fitness of different parasite alleles (36). We have previously conducted these experiments in media containing normal RPMI. We expect that conducting such experiments in diluted RPMI will result in much more rapid detection of differentially selected alleles. Similar approaches are being used to evaluate the fitness consequences of piggyBac insertions in large scale malaria mutagenesis libraries (37, 38). Again, these assays could be made more stringent by use of diluted media to enhance competition.

Compensatory mutations, that reduce the fitness costs of deleterious resistance alleles, are thought to play a major role in resistance evolution (2, 12). In the case of pyrimethamine resistance, copy number amplification of GTP-cyclohydrolase I – the gene at the base of the folate biosynthesis cycle – compensates for reduced efficiency of dihydrofolate reductase later in this biochemical pathway (39, 40). However, rather few other examples are known in malaria. Magnifying the fitness costs of drug resistance alleles using nutrient limitation can accelerate evolution of compensatory changes in laboratory evolution studies. We are currently conducting evolution experiments using diluted media aimed at evolving compensatory changes following introduction of CRISPR generated drug resistance alleles.

## Acknowledgements

This work was supported by National Institutes of Health (NIH) grant R37 AI048071 (to TJCA). Work at Texas Biomedical Research Institute was conducted in facilities constructed with support from Research Facilities Improvement Program grant C06 RR013556 from the National Center for Research Resources.

## Author contributions

S.N., X.L. and T.J.C.A. designed the experiments. S.N conducted experiments, with assistance from M.M-W and M.F. X.L. performed all the NGS analysis and data curation. S. N. X.L. and T.J.C.A. wrote the original manuscript.

Supplementary material

Table S1. Sequence summary for MiSeq amplicon sequencing

[separate file]

## References

1. Andersson DI, Levin BR. The biological cost of antibiotic resistance. Current opinion in microbiology. 1999;2(5):489–93. Epub 1999/10/06. doi: 10.1016/s1369-5274(99)00005-3. PubMed PMID: 10508723.

2. Durão P, Balbontín R, Gordo I. Evolutionary Mechanisms Shaping the Maintenance of Antibiotic Resistance. Trends in microbiology. 2018;26(8):677–91. Epub 2018/02/15. doi: 10.1016/j.tim.2018.01.005. PubMed PMID: 29439838.

3. Shafik SH, Cobbold SA, Barkat K, Richards SN, Lancaster NS, Llinás M, Hogg SJ, Summers RL, McConville MJ, Martin RE. The natural function of the malaria parasite’s chloroquine resistance transporter. Nature communications. 2020;11(1):3922. Epub 2020/08/09. doi: 10.1038/s41467-020-17781-6. PubMed PMID: 32764664; PMCID: PMC7413254.

4. Birnbaum J, Scharf S, Schmidt S, Jonscher E, Hoeijmakers WAM, Flemming S, Toenhake CG, Schmitt M, Sabitzki R, Bergmann B, Fröhlke U, Mesén-Ramírez P, Blancke Soares A, Herrmann H, Bártfai R, Spielmann T. A Kelch13-defined endocytosis pathway mediates artemisinin resistance in malaria parasites. Science (New York, NY). 2020;367(6473):51–9. Epub 2020/01/04. doi: 10.1126/science.aax4735. PubMed PMID: 31896710.

5. Sirawaraporn W, Sathitkul T, Sirawaraporn R, Yuthavong Y, Santi DV. Antifolate-resistant mutants of Plasmodium falciparum dihydrofolate reductase. Proceedings of the National Academy of Sciences of the United States of America. 1997;94(4):1124–9. Epub 1997/02/18. doi: 10.1073/pnas.94.4.1124. PubMed PMID: 9037017; PMCID: PMC19755.

6. Costanzo MS, Hartl DL. The evolutionary landscape of antifolate resistance in Plasmodium falciparum. Journal of genetics. 2011;90(2):187–90. Epub 2011/08/27. doi: 10.1007/s12041-011-0072-z. PubMed PMID: 21869466; PMCID: PMC3212943.

7. Wang X, Mu J, Li G, Chen P, Guo X, Fu L, Chen L, Su X, Wellems TE. Decreased prevalence of the Plasmodium falciparum chloroquine resistance transporter 76T marker associated with cessation of chloroquine use against P. falciparum malaria in Hainan, People’s Republic of China. The American journal of tropical medicine and hygiene. 2005;72(4):410–4. Epub 2005/04/14. PubMed PMID: 15827277.

8. Kublin JG, Cortese JF, Njunju EM, Mukadam RA, Wirima JJ, Kazembe PN, Djimdé AA, Kouriba B, Taylor TE, Plowe CV. Reemergence of chloroquine-sensitive Plasmodium falciparum malaria after cessation of chloroquine use in Malawi. The Journal of infectious diseases. 2003;187(12):1870–5. Epub 2003/06/07. doi: 10.1086/375419. PubMed PMID: 12792863.

9. Ross LS, Fidock DA. Elucidating Mechanisms of Drug-Resistant Plasmodium falciparum. Cell host & microbe. 2019;26(1):35–47. Epub 2019/07/12. doi: 10.1016/j.chom.2019.06.001. PubMed PMID: 31295423; PMCID: PMC6639022.

10. Nkhoma SC, Nair S, Al-Saai S, Ashley E, McGready R, Phyo AP, Nosten F, Anderson TJ. Population genetic correlates of declining transmission in a human pathogen. Molecular ecology. 2013;22(2):273–85. Epub 2012/11/06. doi: 10.1111/mec.12099. PubMed PMID: 23121253; PMCID: PMC3537863.

11. Cerqueira GC, Cheeseman IH, Schaffner SF, Nair S, McDew-White M, Phyo AP, Ashley EA, Melnikov A, Rogov P, Birren BW, Nosten F, Anderson TJC, Neafsey DE. Longitudinal genomic surveillance of Plasmodium falciparum malaria parasites reveals complex genomic architecture of emerging artemisinin resistance. Genome biology. 2017;18(1):78. Epub 2017/04/30. doi: 10.1186/s13059-017-1204-4. PubMed PMID: 28454557; PMCID: PMC5410087.

12. Björkman J, Nagaev I, Berg OG, Hughes D, Andersson DI. Effects of environment on compensatory mutations to ameliorate costs of antibiotic resistance. Science (New York, NY). 2000;287(5457):1479–82. Epub 2000/02/26. doi: 10.1126/science.287.5457.1479. PubMed PMID: 10688795.

13. Levin BR, Perrot V, Walker N. Compensatory mutations, antibiotic resistance and the population genetics of adaptive evolution in bacteria. Genetics. 2000;154(3):985–97. Epub 2000/04/11. PubMed PMID: 10757748; PMCID: PMC1460977.

14. Hartl DL, Clark AG, Sinauer A. Principles of population genetics. Sunderland: Sinauer Associates, Inc. Publishers; 2018.

15. Duvalsaint M, Conrad MD, Tukwasibwe S, Tumwebaze PK, Legac J, Cooper RA, Rosenthal PJ. Balanced impacts of fitness and drug pressure on the evolution of PfMDR1 polymorphisms in Plasmodium falciparum. Malaria journal. 2021;20(1):292. Epub 2021/07/02. doi: 10.1186/s12936-021-03823-x. PubMed PMID: 34193148; PMCID: PMC8247092.

16. Gabryszewski SJ, Dhingra SK, Combrinck JM, Lewis IA, Callaghan PS, Hassett MR, Siriwardana A, Henrich PP, Lee AH, Gnädig NF, Musset L, Llinás M, Egan TJ, Roepe PD, Fidock DA. Evolution of Fitness Cost-Neutral Mutant PfCRT Conferring P. falciparum 4-Aminoquinoline Drug Resistance Is Accompanied by Altered Parasite Metabolism and Digestive Vacuole Physiology. PLoS pathogens. 2016;12(11):e1005976. Epub 2016/11/11. doi: 10.1371/journal.ppat.1005976. PubMed PMID: 27832198; PMCID: PMC5104409.

17. Ross LS, Dhingra SK, Mok S, Yeo T, Wicht KJ, Kümpornsin K, Takala-Harrison S, Witkowski B, Fairhurst RM, Ariey F, Menard D, Fidock DA. Emerging Southeast Asian PfCRT mutations confer Plasmodium falciparum resistance to the first-line antimalarial piperaquine. Nature communications. 2018;9(1):3314. Epub 2018/08/18. doi: 10.1038/s41467-018-05652-0. PubMed PMID: 30115924; PMCID: PMC6095916.

18. Straimer J, Gnädig NF, Stokes BH, Ehrenberger M, Crane AA, Fidock DA. Plasmodium falciparum K13 Mutations Differentially Impact Ozonide Susceptibility and Parasite Fitness In Vitro. mBio. 2017;8(2). Epub 2017/04/13. doi: 10.1128/mBio.00172-17. PubMed PMID: 28400526; PMCID: PMC5388803.

19. Nair S, Li X, Arya GA, McDew-White M, Ferrari M, Nosten F, Anderson TJC. Fitness Costs and the Rapid Spread of kelch13-C580Y Substitutions Conferring Artemisinin Resistance. Antimicrobial agents and chemotherapy. 2018;62(9). Epub 2018/06/20. doi: 10.1128/aac.00605-18. PubMed PMID: 29914963; PMCID: PMC6125530.

20. Tilman D. Resource competition and community structure. Monographs in population biology. 1982;17:1–296. Epub 1982/01/01. PubMed PMID: 7162524.

21. Wale N, Sim DG, Jones MJ, Salathe R, Day T, Read AF. Resource limitation prevents the emergence of drug resistance by intensifying within-host competition. Proceedings of the National Academy of Sciences of the United States of America. 2017;114(52):13774–9. Epub 2017/12/14. doi: 10.1073/pnas.1715874115. PubMed PMID: 29233945; PMCID: PMC5748215.

22. Wale N, Sim DG, Read AF. A nutrient mediates intraspecific competition between rodent malaria parasites in vivo. Proceedings Biological sciences. 2017;284(1859). Epub 2017/07/28. doi: 10.1098/rspb.2017.1067. PubMed PMID: 28747479; PMCID: PMC5543226.

23. Letten AD, Hall AR, Levine JM. Using ecological coexistence theory to understand antibiotic resistance and microbial competition. Nature ecology & evolution. 2021;5(4):431–41. Epub 2021/02/03. doi: 10.1038/s41559-020-01385-w. PubMed PMID: 33526890.

24. Paulander W, Maisnier-Patin S, Andersson DI. The fitness cost of streptomycin resistance depends on rpsL mutation, carbon source and RpoS (sigmaS). Genetics. 2009;183(2):539–46, 1si–2si. Epub 2009/08/05. doi: 10.1534/genetics.109.106104. PubMed PMID: 19652179; PMCID: PMC2766315.

25. Petersen A, Aarestrup FM, Olsen JE. The in vitro fitness cost of antimicrobial resistance in Escherichia coli varies with the growth conditions. FEMS microbiology letters. 2009;299(1):53–9. Epub 2009/08/22. doi: 10.1111/j.1574-6968.2009.01734.x. PubMed PMID: 19694815.

26. Bunditvorapoom D, Kochakarn T, Kotanan N, Modchang C, Kümpornsin K, Loesbanluechai D, Krasae T, Cui L, Chotivanich K, White NJ, Wilairat P, Miotto O, Chookajorn T. Fitness Loss under Amino Acid Starvation in Artemisinin-Resistant Plasmodium falciparum Isolates from Cambodia. Scientific reports. 2018;8(1):12622. Epub 2018/08/24. doi: 10.1038/s41598-018-30593-5. PubMed PMID: 30135481; PMCID: PMC6105667.

27. LeRoux M, Lakshmanan V, Daily JP. Plasmodium falciparum biology: analysis of in vitro versus in vivo growth conditions. Trends in parasitology. 2009;25(10):474–81. Epub 2009/09/15. doi: 10.1016/j.pt.2009.07.005. PubMed PMID: 19747879.

28. Desai SA. Insights gained from P. falciparum cultivation in modified media. TheScientificWorldJournal. 2013;2013:363505. Epub 2013/08/21. doi: 10.1155/2013/363505. PubMed PMID: 23956690; PMCID: PMC3727134.

29. Brown AC, Guler JL. From Circulation to Cultivation: Plasmodium In Vivo versus In Vitro. Trends in parasitology. 2020;36(11):914–26. Epub 2020/09/23. doi: 10.1016/j.pt.2020.08.008. PubMed PMID: 32958385.

30. Liu J, Istvan ES, Gluzman IY, Gross J, Goldberg DE. Plasmodium falciparum ensures its amino acid supply with multiple acquisition pathways and redundant proteolytic enzyme systems. Proceedings of the National Academy of Sciences of the United States of America. 2006;103(23):8840–5. Epub 2006/05/30. doi: 10.1073/pnas.0601876103. PubMed PMID: 16731623; PMCID: PMC1470969.

31. Istvan ES, Dharia NV, Bopp SE, Gluzman I, Winzeler EA, Goldberg DE. Validation of isoleucine utilization targets in Plasmodium falciparum. Proceedings of the National Academy of Sciences of the United States of America. 2011;108(4):1627–32. Epub 2011/01/06. doi: 10.1073/pnas.1011560108. PubMed PMID: 21205898; PMCID: PMC3029723.

32. Ebel E, Kuypers F, Lin C, Petrov D, Egan E. Common host variation drives malaria parasite fitness in healthy human red cells. bioRxiv; 2020.

33. Cook RD. Detection of Influential Observation in Linear Regression. Technometrics. 1977;19(1):15–8. doi: 10.1080/00401706.1977.10489493.

34. Müller IB, Hyde JE. Folate metabolism in human malaria parasites--75 years on. Molecular and biochemical parasitology. 2013;188(1):63–77. Epub 2013/03/19. doi: 10.1016/j.molbiopara.2013.02.008. PubMed PMID: 23500968.

35. Kicska GA, Ting LM, Schramm VL, Kim K. Effect of dietary p-aminobenzoic acid on murine Plasmodium yoelii infection. The Journal of infectious diseases. 2003;188(11):1776–81. Epub 2003/11/26. doi: 10.1086/379373. PubMed PMID: 14639551.

36. Li X, Kumar S, McDew-White M, Haile M, Cheeseman IH, Emrich S, Button-Simons K, Nosten F, Kappe SHI, Ferdig MT, Anderson TJC, Vaughan AM. Genetic mapping of fitness determinants across the malaria parasite Plasmodium falciparum life cycle. PLoS genetics. 2019;15(10):e1008453. Epub 2019/10/15. doi: 10.1371/journal.pgen.1008453. PubMed PMID: 31609965; PMCID: PMC6821138.

37. Zhang M, Wang C, Otto TD, Oberstaller J, Liao X, Adapa SR, Udenze K, Bronner IF, Casandra D, Mayho M, Brown J, Li S, Swanson J, Rayner JC, Jiang RHY, Adams JH. Uncovering the essential genes of the human malaria parasite Plasmodium falciparum by saturation mutagenesis. Science (New York, NY). 2018;360(6388). Epub 2018/05/05. doi: 10.1126/science.aap7847. PubMed PMID: 29724925; PMCID: PMC6360947.

38. Bronner IF, Otto TD, Zhang M, Udenze K, Wang C, Quail MA, Jiang RH, Adams JH, Rayner JC. Quantitative insertion-site sequencing (QIseq) for high throughput phenotyping of transposon mutants. Genome research. 2016;26(7):980–9. Epub 2016/05/20. doi: 10.1101/gr.200279.115. PubMed PMID: 27197223; PMCID: PMC4937560.

39. Kümpornsin K, Modchang C, Heinberg A, Ekland EH, Jirawatcharadech P, Chobson P, Suwanakitti N, Chaotheing S, Wilairat P, Deitsch KW, Kamchonwongpaisan S, Fidock DA, Kirkman LA, Yuthavong Y, Chookajorn T. Origin of robustness in generating drug-resistant malaria parasites. Molecular biology and evolution. 2014;31(7):1649–60. Epub 2014/04/18. doi: 10.1093/molbev/msu140. PubMed PMID: 24739308; PMCID: PMC4069624.

40. Nair S, Miller B, Barends M, Jaidee A, Patel J, Mayxay M, Newton P, Nosten F, Ferdig MT, Anderson TJ. Adaptive copy number evolution in malaria parasites. PLoS genetics. 2008;4(10):e1000243. Epub 2008/11/01. doi: 10.1371/journal.pgen.1000243. PubMed PMID: 18974876; PMCID: PMC2570623.

